# A paradigm for skilled forelimb reaching by head-restrained mice

**DOI:** 10.64898/2026.05.25.727681

**Authors:** Annabel Chang, Adithya Madduri, Janet Berrios Wallace, Junzheng Wang, Kimberly Reinhold

## Abstract

Forelimb reaching tasks are widely used in neuroscience across species. In mice, pellet reaching is especially useful because it requires fine motor control. However, few automated pellet-delivery systems are well suited for head-restrained reaching, where pellet position must be highly consistent. Here we present a complete pipeline for head-restrained, or head-fixed, forelimb reaching in mice. The pipeline includes a simple customizable pellet-presentation rig, an effective training protocol, and analysis code to detect reaches and classify their outcomes. To demonstrate pipeline utility, we quantified interruption of an ongoing reach. The temporal requirement for initiating cancellation was dependent on the phase of the reach, lengthening significantly as the reach neared completion. These results show that the pipeline can resolve fine phase-dependent features of skilled behavior and provide a practical framework for studying motor control in head-restrained mice.

## Introduction

Paradigms for skilled forelimb behavior in rodents are valuable because they allow motor initiation, learning, and control to be studied across many levels of the nervous system^1–6^. Reaching for food pellets is a common example^7–12^. Although this behavior is often studied in freely moving mice, head-fixed reaching is increasingly being used in neuroscience^13–20^. Head fixation constrains posture and reduces the animal’s behavioral repertoire. However, it can also limit competing behaviors, allowing the task to focus on repeated execution of the trained reach. Moreover, head fixation is compatible with fixed-position equipment such as microscopes and electrodes^21,22^. For these reasons, we developed a complete pipeline for head-fixed forelimb reaching in mice.

The pipeline has three parts: a motorized pellet-delivery apparatus, a training protocol, and a video-based analysis pipeline. Together, these tools provide a practical and transparent resource for laboratories seeking to implement this task.

To study reaching toward a learned location, food pellets must be delivered to a consistent position in front of the mouse. While various pellet dispensers exist, few are optimized for the precise and repeatable pellet placement required in head-fixed reaching^23–31^. We therefore developed a simple system using two stepper motors: one to dispense pellets and one to position them in front of the animal. The device is controlled by programmable Arduino microcontrollers and built from custom laser-cut or 3D-printed components. It was designed to be inexpensive, reproducible, and easy to assemble.

Training is also a challenge in head-fixed reaching. In freely moving tasks, mice can practice reaches at will, but head fixation restricts the time available for daily training and may suppress spontaneous reaching. Indeed, most mice do not readily reach for pellets when first head-fixed. We therefore developed a shaping protocol to train mice to perform the behavior reliably under these conditions.

A final critical step is accurate measurement of behavior. Rather than using beam breaks or touch sensors, we relied on video as the sole readout. Video-based analysis is noninvasive, avoids electrical noise, and captures the full reach trajectory rather than a single event. It also allows direct confirmation of reach outcome, such as whether the pellet was successfully retrieved and consumed. We developed an expert-system pipeline to detect reaches and classify outcomes. This approach is computationally simple and interpretable, although it requires some manual parameter tuning.

To illustrate the utility of this paradigm, we examined cancellation of an ongoing reach. We considered two simple possibilities: that cancellation time might be fixed across the movement, or that it might depend on the current reach phase. We found that the cancellation time depended on reach phase, showing that the behavioral pipeline is sensitive enough to resolve fine features of ongoing skilled behavior.

## Materials and Methods

### Animals

All procedures were approved by the President and Fellows of Harvard College Institutional Animal Care and Use Committee under protocol #IS00000571-6. Experiments were performed in male and female mice older than 3 months. Females were group-housed, and males were singly housed. Mice were housed on a reverse light cycle. Ambient room temperature was maintained at 75 °F, with 45% relative humidity. During behavioral training, mice were food restricted to approximately 85% of baseline body weight to maintain motivation, with daily monitoring of health and weight.

### Materials

See **Table 1** and Code and Materials Availability statement.

**Table 1.** Materials for pellet loading and presentation, excluding custom components (for custom designs, see Code and Materials Availability).

| Part name | Quantity needed | Areas used | Cost (USD) | Unit price |
| --- | --- | --- | --- | --- |
| 324 NEMA-17 Stepper Motor | 2 | Positioner ensemble and loading ensemble | 28 | 14 |
| 18-8 Stainless Steel Hex Drive Flat Head Screw, M3 x 0.5 mm Thread, 50 mm Long | 8 | Positioner ensemble and loading ensemble | 20.66 | 2.58 |
| XYZ Axis Manual Precision Linear Stage 40x40mm Trimming Bearing Tuning Platform Sliding Table (Bearing Number: LD40-LM, Bearing Type: Roller Bearing, UPC: 709046491161, ASIN: B07D7N9GT6) | 1 | Pellet stage ensemble | 125 | 125 |
| MB4 - Aluminum Breadboard 4" x 6" x 1/2", 1/4"-20 Taps | 1 | Pellet stage ensemble | 52.34 | 52.34 |
| SR4 - Compact Cage Assembly Rod, 4" Long, Ø4 mm | 2 | Pellet stage ensemble | 21.5 | 10.75 |
| 1/8th and 1/16th inch-thick Acrylic Sheets | 2 each | Pellet stage and positioner ensembles | 79.68 | 19.92 |
|  |  | **Total cost** | **327.18** |  |

## Methods: Behavior Rig

### Overview

We built a behavior rig to automatically and precisely position food pellets (Bio-Serv F05301) in front of the mouse (**Fig. 1A**). The rig consisted of three main components: (1) a “mouse house”: a stable platform where the head-fixed mouse sat and rested its paws (on the edge of the platform called the “perch”); (2) a “pellet positioner”: a motorized disk used to move individual pellets to a consistent, precise spot in front of the mouse; and (3) a “pellet loader”: a high-capacity hopper that stored and dispensed all the pellets needed for a full training session (typically over one hour).

**Figure 1.**
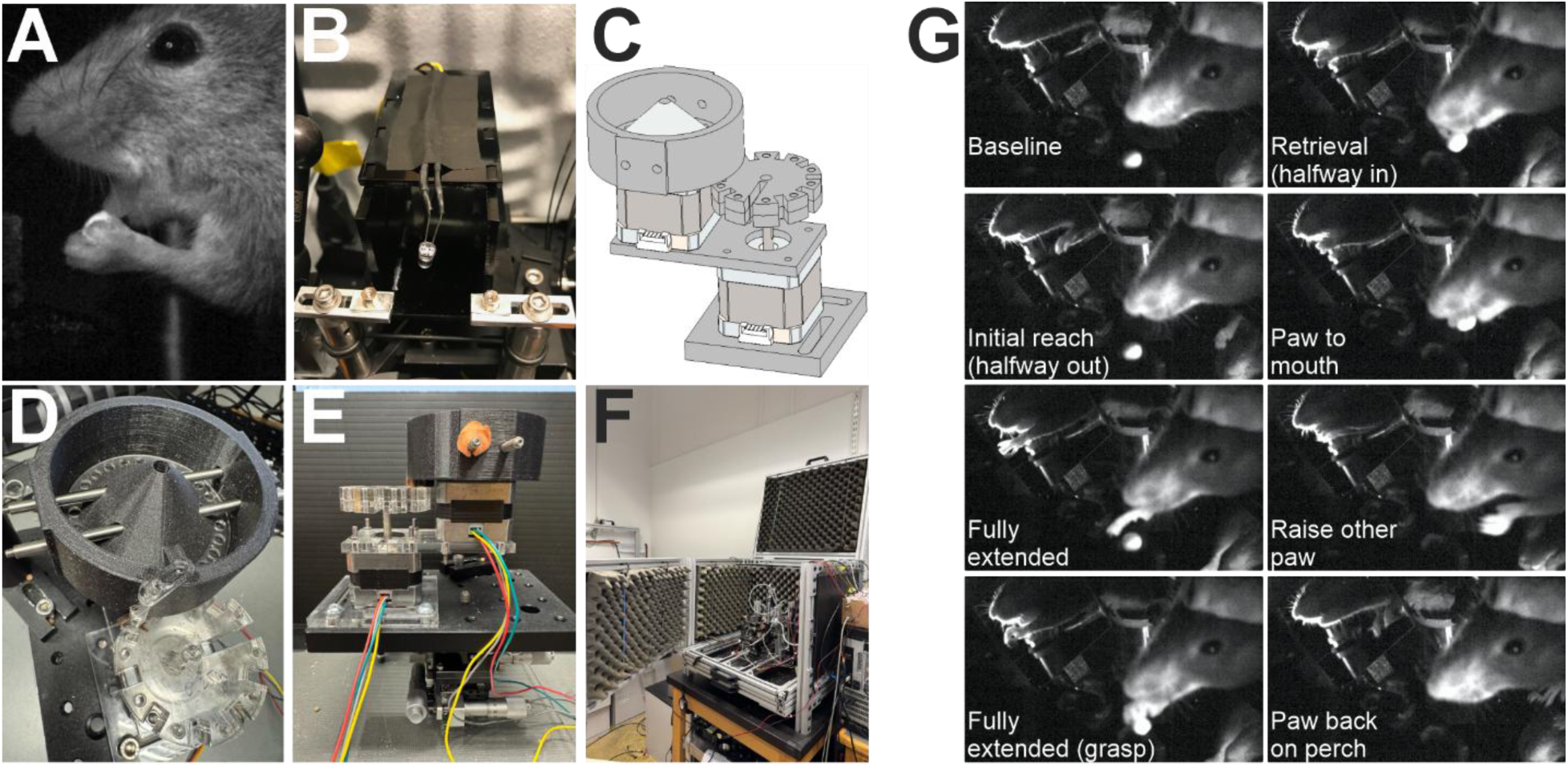
Rig design for head-fixed mouse reaching behavioral paradigm. **(A)** Example video frame showing an intermediate reach stage as mouse learns to reach forward, grasp the food pellet, flip the paw and raise the pellet to the mouth. **(B)** Mouse house. (**C)** CAD assembly for pellet positioner and pellet loader. **(D)** Top-down picture of pellet loader (top, black) and pellet positioner (bottom, clear). **(E)** Side-view picture of pellet loader and pellet positioner. **(F)** Enclosure box with sound-proofing foam. **(G)** Phases of a typical reach in temporal order from left column top to bottom, then right column top to bottom.

The mouse house (**Fig. 1B**) was a platform with two walls and a small, removable roof made from laser-cut acrylic. The walls and ceiling were provided to help the mouse feel enclosed and therefore more comfortable in the apparatus. Thorlabs posts flanking the platform enabled head-fixation.

The pellet positioner was a flat disk mounted to the axle of a stepper motor. The disk had recessed circles around the edge (**Fig. 1C,D,E**). Single pellets sat in these circles. As the motor turned, the disk rotated, and the pellets moved, bringing one pellet at a time directly in front of the mouse’s paw. The remaining pellets on the disk were spaced far enough apart that the mouse could not reach them while head-fixed. At each turn of the motor, the currently available pellet was replaced by the next pellet. After pellets moved past the mouse, they were dislodged from the disk by a paintbrush secured to the pellet loader (described next).

Like the pellet positioner, the pellet loader (i.e., food hopper) consisted of a laser-cut flat disk mounted to the axle of a stepper motor (**Fig. 1C,D,E**). Numerous small holes lined the perimeter of this disk. Pellets fell into these holes and rotated around the central axle. The disk was housed within a larger, 3D-printed container. Although this container held many more pellets than the total number of holes, the disk was thin, so that only one pellet could occupy each hole. The other pellets sat piled above. The disk rotated on a platform featuring a single opening located beneath a 3D-printed lip. Individual pellets fell through this opening onto the pellet positioner, and the lip prevented more than one pellet from being dispensed at a time.

Both the pellet positioner and the pellet loader were jointly mounted onto a single laser-cut platform. This ensured consistent relative positioning of these two components. This platform was then attached to a 3D positioner/micrometer, enabling 3-axis positioning of the pellet with respect to the mouse paw. The mouse house was not coupled to this positioner but was separately mounted on the behavior frame. Consequently, the food pellet position could be controlled independently of the mouse’s position.

We chose this design using two stepper motors for simplicity. However, because the system lacked real-time feedback to adjust the motor angles, any initial angular misalignments persisted throughout a training session. Hence, it was critical to prevent initial misalignment of the pellet positioner. To ensure precision, we incorporated a fixed-position cutout beneath the pellet positioner wheel to serve as a visual guide, allowing the experimenter to correctly calibrate the initial angle of the pellet positioner’s stepper motor. Any jamming that resulted in an angular error during a session could be fixed by disconnecting the Arduino powering the stepper motor, manually resetting the angle, and resuming the session. Such malfunctions were quite infrequent (<1 per week given 6 training sessions per day).

### Electronic Control

Both stepper motors (for the pellet positioner and pellet loader) were controlled by an Arduino V2 Motor Shield connected to an Arduino Uno. The Arduino Uno also provided output signals to manage other behavioral components, such as LEDs for visual cues or optogenetic stimulation. Behavioral events were recorded onto a microSD card via an Arduino Ethernet shield. Thus, the electronic control of the behavior system was both simple and flexible.

### Frame and Behavior Enclosure Box

Mice performed the behavior in complete darkness to enforce reaching that was memory-guided rather than visually guided. To achieve darkness and exclude ambient light, the entire behavior rig was enclosed in a larger chamber (**Fig. 1F**) custom-built from T-slotted aluminum extrusions (the frame) and black cardboard (the walls). On the inside, the walls were lined with sound-proofing foam. We included a drawer at the bottom of the enclosure box to collect fallen pellets to make cleaning easier.

### Low-Speed Video Acquisition

We positioned a low-speed camera (30 frames per second) to the side of the mouse to capture continuously recorded video of the mouse reaching and consuming food pellets. The video data was transferred to a low-cost DVR that logged frames onto a microSD card. Following each training session, we transferred the video files from the microSD card to a central computer. This configuration allowed multiple behavior rigs to operate in parallel without requiring a dedicated computer for each rig. Although the low-cost DVR occasionally dropped frames, this did not disrupt our custom analysis pipeline, as all behavioral events were detected within the video’s own temporal reference frame and realigned post hoc to the Arduino commands (see Temporal Synchronization and IR Signaling below).

### High-Speed Paw Tracking

For experiments requiring high-speed paw tracking, we added a second high-speed Flea3 FLIR camera, and we positioned two perpendicular mirrors below and to the side of the forelimb. The angled Flea3 camera was mounted above the mouse and directed toward the mirrors to capture the forelimb from multiple perspectives (**Fig. 1G**). Consequently, a single high-speed camera frame provided three simultaneous views of the mouse’s paw: from below, from the side, and from above. High-speed video was acquired at 256 frames per second (fps) using FlyCapture software and streamed directly to a computer. The reach trajectory could be tracked by DeepLabCut^32,33^ and reconstructed in 3D based on the perpendicular mirrors’ views. High-speed tracking was not implemented on every rig, but when used, it was essential to angle the mirrors to prevent flashing lights–such as cues or optogenetic stimuli–from reflecting into the low-speed camera’s view of the reach zone. This was because light artifacts might interfere with the automated reach detection based on the video.

### Infrared (IR) Illumination

The low-speed camera included a built-in array of infrared (IR) LEDs. This array of IR LEDs was required to provide illumination of the mouse for video imaging. Because mice do not perceive IR light, the animals remained functionally in the dark^34^.

### Acoustic Masking of Motor Noise

The stepper motor generated noise, an audible mechanical sound. The mice could potentially use this sound as a cue to reach. To mitigate this, we implemented several controls. First, we significantly degraded the utility of the motor sound as a reliable predictor of the food pellet. We skipped loading pellets into many of the slots on the pellet positioner. Consequently, the disk rotated frequently without presenting a pellet (i.e., moved an empty slot toward the mouse). Second, we reduced the salience of the motor sound by playing a masking sound. This masking sound was a continuous recording of the stepper motor spinning. We played this masking sound through two speakers positioned on either side of the stepper motor. This acted as a potent, although still imperfect, masker of the pellet positioner’s movement. A small fraction of the mice still attempted to use the motor sound as a cue during learning (<15%).

### Olfactory Masking and CPU Fan

An important control was to prevent the mouse from using olfactory cues to detect the presence or proximity of the food pellet. To achieve this, we used a two-part strategy. First, we positioned a CPU fan to direct a stream of air toward the mouse’s face. This created air turbulence, preventing the mouse from spatially localizing the food pellet. Second, we positioned a pile of food pellets near the mouse but out of reach. This pile of pellets served as an olfactory masker.

## Methods: Mouse Training Protocol

### Food Restriction and Handling (2 days)

To ensure motivation for the task, we placed mice on a food restriction schedule. We presented food in the home cage in the form of Bio-Serv pellets and peanut butter to familiarize mice with these foods. During this period, we also habituated the animals to experimenter contact to minimize stress. On Day 1, we placed the experimenter’s gloved hand in the corner of the home cage, allowing the mouse to investigate voluntarily. On Day 2, we introduced a palatable reward (peanut butter) on a finger to encourage direct interaction. On Day 3, we transitioned to lifting the mouse in the palm of the experimenter’s hand. Mice learned to be transported in the palm of the hand without discomfort or anxiety.

### Rig Habituation and Licking (3 days)

Initial training occurred on a behavior rig situated *outside* of the enclosure box, so that the experimenter could interact directly with the mouse. We removed the pellet loader and electronically disconnected the motors for this “training rig”. Only the mouse house and pellet positioner were present. The experimenter could manually turn the pellet positioner disk to simulate its movement in the automated rig. Initial days on this training rig focused on associating the head-restrained posture with reward delivery. We provided peanut butter (**Fig. 2A**) or food pellets (**Fig. 2B**) directly to the mouth. For the first day, we positioned the peanut butter or food pellets on either the surface of the pellet positioner disk or a small platform. We raised the surface to mouth-level, so the mouse could access the food by licking. This platform or the pellet positioner disk served as a stationary surface attached to a micrometer enabling precise up-down adjustments to position the pellets very near the animal’s mouth. Importantly, many pellets covered this surface, ensuring a high success rate when the mouse attempted to access a pellet with the tongue. Occasionally, mice refused whole pellets due to the mechanical unfamiliarity of chewing while head-fixed. In this case, we crushed the pellets into smaller pieces. It was important to encourage eating pellets, not just licking peanut butter. To do this, as soon as the mice were comfortable, we switched out the peanut butter for a dense layer of pellets that could be easily accessed by the tongue and mouth. The goal of this phase was to achieve 30 minutes of consistent pellet consumption without anxiety. Once mice were comfortable eating pellets off of the platform/positioner disk, we began to offer individual food pellets “glued” by peanut butter to the end of a stick (Q-tip wooden end). This “pellet-on-a-stick” method served as the primary food delivery mechanism for several of the next training steps, as indicated below. To present the food pellet to the mouse, we offered the pellet just below the mouth. Mice learned to retrieve this pellet by licking. Care was taken to avoid pressing the pellet against the animal’s face. Some mice were more comfortable eating pieces of crushed-up pellet rather than the whole pellet at this stage.

**Figure 2.**
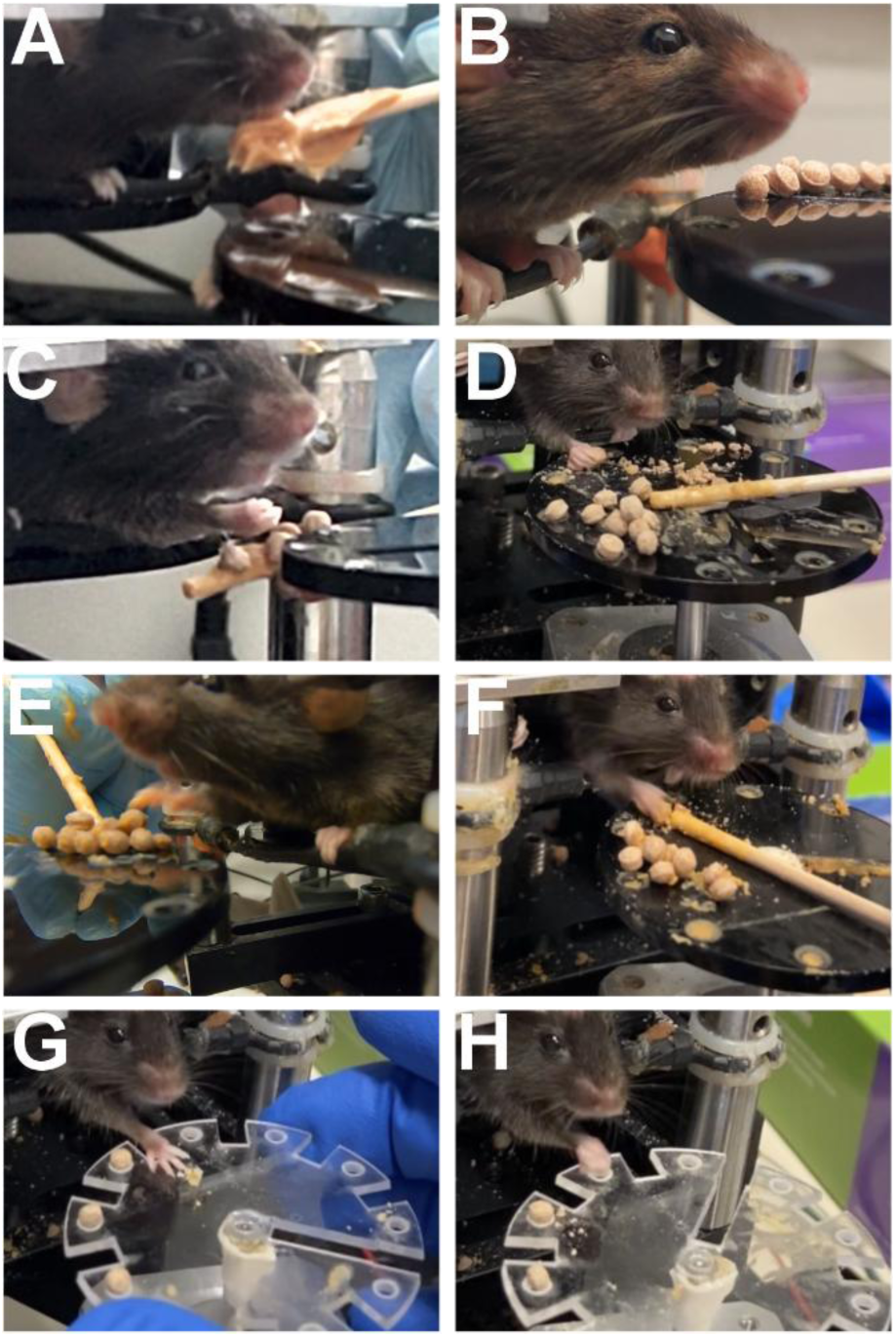
Example stages of reach training. **(A)** Licking peanut butter off stick. **(B)** Pellets positioned below the mouth, enabling spontaneous licking to obtain food. **(C)** Reach to touch stick. **(D)** Reach to pellets on platform. **(E)** Reach to pellet clumps on platform. **(F)** Stick orientation to discourage laterally swatting the stick far away from the pellet. **(G)** Spontaneously initiated reach but incorrect pellet targeting. **(H)** Spontaneously initiated reach with successful targeting.

### “Pellet-on-a-Stick” Part 1: Shaping the Reaching Trajectory (1-3 days)

We trained the reaching behavior by shaping. Initially, we induced forelimb movement by applying a tactile cue—gently tapping the mouse’s shoulder, face or whiskers with the wooden stick. This elicited a swatting motion toward the stick. We reinforced any contact between the right paw and the target stick with a reward, i.e., presentation of a food pellet “glued” to the end of a different stick with peanut butter (**Fig. 2C**). As the mouse began to swat reliably, we adjusted the position of the target stick outward to require full limb extension.

### Notes on Shaping the Reaching Trajectory

Some mice did not respond to the tactile cue. Two other strategies were possible to elicit contact between the mouse’s paw and the stick. First, for mice that fidgeted frequently, placing the stick near the animal’s paw sometimes resulted in the mouse placing the paw spontaneously onto the stick. When immediately rewarded, this behavior was repeated. Second, some mice reached directly to the stick when the food pellet or peanut butter was held beneath the nose for a few minutes. In general, mice preferred peanut butter over food pellets, so we were careful to use as little peanut butter as possible to avoid devaluing the food pellets relative to the peanut butter.

### Engaging the Forelimb Early in Training

It was critical to engage the mouse’s forelimb early in training. Extending the Rig Habituation and Licking phase for too many days led to animals becoming highly proficient at licking the pellet without any forelimb engagement. Once these individuals over-trained on the licking behavior, they stubbornly persisted in licking rather than attempting to reach, even after the pellet was moved out of reach. This is worth emphasizing again: in this step and subsequent steps, it was vital to prevent mice from successfully obtaining pellets using the tongue without engaging the forelimb.

### Many Accessible Pellets Early in Training

It was helpful to position a layer of pellets on the platform/positioner disk beneath the stick—hence, whenever a mouse swiped for the stick and missed, the paw made tactile contact with the food. For some animals, simply recognizing that food pellets were available was sufficient to immediately trigger the full reaching motion with clear volitional intent. In contrast, other animals seemed unable to plan the motor sequence of the reach, even after understanding the availability of within-reach pellets.

### “Pellet-on-a-Stick” Part 2: Grasping and Retrieval (1-4 days)

To transition from a swat to a functional reach, we next prioritized the mechanics of grasping and supination (the rotation of the paw to orient the palm toward the mouth). The goal was to get the mouse grasping and attempting to retrieve the pellet. There were several sub-steps. First, when the mouse touched the pellet, we manually guided the paw toward the mouth to mimic successful retrieval. Second, we required the mouse to push the pellet off the stick and into its mouth. Third, the pellet was moved progressively further from the animal’s mouth to encourage the active grasping and manipulation required to transfer pellets from the delivery stick to the mouth. As proficiency increased, we faded the manual assistance and moved the target pellet onto the platform/positioner disk, requiring the mouse to pluck the pellet directly from this surface (**Fig. 2D**).

### Spontaneous Reaching

Some animals (<20%) were able to bypass the “pellet-on-a-stick” training phase entirely. During early training stages, these individuals tended to spontaneously attempt to push or manipulate pellets into the mouth. When this behavior emerged, we gradually moved the platform/positioner disk with pellets farther from the animal to encourage reaching. Typically, these mice became proficient at the full reaching sequence—including grasping and lifting the pellet to the mouth—within two to three days.

### Reaching for Pellet Clumps

Some mice initially struggled to manipulate the small food pellets but exhibited spontaneous reaches. To assist these individuals, we offered larger clumps of pellets bonded with peanut butter (**Fig. 2E**). These clumps were easier to push toward the mouth, bypassing the need for fine motor control while mice continued to develop proficiency handling the small pellets.

### Transitioning from Swatting the Stick to Handling the Pellet

For mice requiring the “pellet-on-a-stick” phase, it was essential to quickly transition the reward criterion from simple swatting of the stick to active handling of the pellet. If animals became too accustomed to knocking the pellet off the stick to receive a reward, they often struggled to attempt the more complex movements required for pellet grasping and retrieval. To encourage the correct motor behavior, we adjusted the orientation of the delivery stick according to the specific needs of each animal.

### Downward Angle

The stick was angled higher at its lateral end and sloped downward toward the mouth medially, so that the paw naturally slid toward the face upon contact.

### Anterior-Posterior Alignment

The stick was positioned parallel to the mouse’s midline (**Fig. 2F**), forcing the animal to make contact at the tip (closest to the mouth). This prevented lateral reaching, where a mouse would grasp a lateral part of the stick far away from the pellet.

### Notes on Grasping and Retrieval

The grasp and supination in the head-fixed posture could be tricky to elicit. Hence, we initially rewarded any rotation of the paw toward the face, even if the pellet was not successfully transferred to the mouth.

### Reaching with the Right or Left Forelimb

We trained all mice to reach with the right-side forelimb. To accomplish this, we strictly avoided rewarding reaches initiated with the non-target forelimb (e.g., using the left limb when the right was required).

### Consistency

Progress varied significantly between individuals. Some mice mastered multiple steps in a single day’s session and others required several days for a single transition. Importantly, we maintained a strict training progression and avoided “backtracking” (e.g., letting the animal lick for pellets during a reaching stage).

### “Pellet-on-a-Stick” Part 3: From Manually Cued to Self-Initiated Reaches (1-3 days)

Once mice consistently executed the reach-to-grasp sequence in response to a shoulder poke, we transitioned to more distal sensory cues. First, we replaced the shoulder poke with a whisker cue, where the delivery stick was used to lightly brush the whiskers on the side of the face ipsilateral to the reaching arm. The touch point was moved incrementally from the shoulder toward the whiskers over several trials. We then moved the delivery stick farther away to the level of the platform/positioner disk. Animals could visually detect the stick, triggering reaching. Eventually, we removed the stick entirely, and mice began to reach spontaneously toward pellets resting on the platform/positioner disk.

### Reach Refinement and Turning Disk Training (1-3 days)

In the final stage of reach training, we refined the self-initiated reach. At the beginning of this stage, mice attempted to grasp pellets from the surface of the pellet positioner disk. We initially densely loaded the disk with many pellets to increase the chances of grasping a pellet. If mice struggled to successfully retrieve a single pellet, we created larger chunks of pellets stuck together with peanut butter. Often, animals were able to successfully push these larger chunks into the mouth, even before the mice were capable of manipulating individual pellets. Once animals became proficient at manipulating chunks or single pellets, we reduced the density of the pellets. We began to present pellets in a single consistent location (**Fig. 2G**). Lastly, to simulate the conditions of the automated rig, we manually turned the pellet positioner disk to present pellets repeatedly in the same place. To encourage outward extension of the forelimb, we moved the pellets away from the mouse to the maximum reachable distance (**Fig. 2H**).

### Transition to Automated Task (1-2 days)

Upon mastering the mechanics of the reach, mice were transferred to the automated behavior rig in the enclosure box. Starting at 80-100% of the pellet positioner slots loaded with pellets, over several training sessions, we gradually reduced the frequency of pellets loaded in the pellet positioner slots to 30-40%. This ensured that initial reaches in the enclosure box were successful (pellet always available), reinforcing the reaching behavior in the new context of the automated rig. However, we quickly reduced pellet availability to only the time window after the cue. This was required for mice to learn a cue-pellet pairing. This successfully transitioned the mice from continuous reinforcement to the trial-based structure of the final paradigm, where reaches were triggered by specific cues.

### Notes on Transition to Automated Task

Most mice successfully reached without prompting in the new environment of the automated rig within 5-10 minutes. However, although the training rig was nearly identical to the automated rig (minus the enclosure box), some mice required further prompting to reach after being transitioned off of the training rig onto the automated rig. Two approaches were used to prompt reaching in these mice. First, we used the olfactory cue of peanut butter to prompt appetitive behavior. We placed a small smear of peanut butter on the pellet loader in front of the mouse. After a few minutes, this prompted the mouse to attempt reaching forward to obtain food. Second, in some cases, we reintroduced the stick near the mouse’s face, as a learned cue to trigger reaching. The goal was simply to trigger a reach. Once mice reached forward and discovered the accessible pellet, they typically continued to reach. Another challenge in some cases was a mouse reaching to the wrong location and consistently failing to retrieve a pellet. Despite the reproducible design of the training and automated rigs, some mice changed their egocentric reach targets after being transferred to the automated rig. In this case, using the micrometers/positioners on the pellet presentation apparatus was helpful to position the food pellet at the terminus of the mouse reach. Once animals began to retrieve food reliably, it was possible to further shape the reaching trajectory, if necessary. Another option, in mice that were well-trained to reach to the tip of the stick, was to use the stick tip as a cue to guide the reach toward the pellet’s location. After mice became comfortable reaching, they performed the behavior quite reliably. As a note, if a trained animal suddenly stopped reaching for >10 min in the middle of a normal behavior session, this was typically because the animal had urinated. In this case, the mouse was fed some peanut butter and immediately returned to the home cage.

## Methods: Video Analysis Pipeline

### Overview

The video analysis pipeline identified and monitored specific Regions of Interest (ROIs), or “zones”, within the video frames to classify behavioral events. By tracking pixel-intensity signals within these predefined ROIs—such as the region traversed by the paw during the reach, the pellet loading zone, or the mouth—the system detected events including reaching, pellet retrieval, and chewing over time. The pipeline consisted of four primary stages: 1. ROI Definition: Identifying behaviorally relevant spatial zones within the video frames. 2. Signal Extraction: Gathering temporal signals (e.g., pixel intensity or motion) from these ROIs across all frames of the video. 3. Heuristic Signal Processing: Employing rule-based algorithms to detect discrete behavioral events, such as reaches, pellet removals, chewing, and LED cues. 4. Outcome Refinement: Discriminating between reach outcomes, specifically distinguishing successful pellet retrievals from attempts when the mouse dropped the pellet.

### Definition of Behavior-Relevant Regions of Interest (ROIs)

Identifying behaviorally relevant spatial zones within the video frames was simplified by the head-fixed preparation, which ensured consistent positioning of the mouse’s head and the food pellet across a training session. We defined several distinct Regions of Interest (ROIs) (**Fig. 3**): the reach, pellet, eat, perch, and lick zones, alongside task-specific ROIs for cues and distractors.

**Figure 3.**
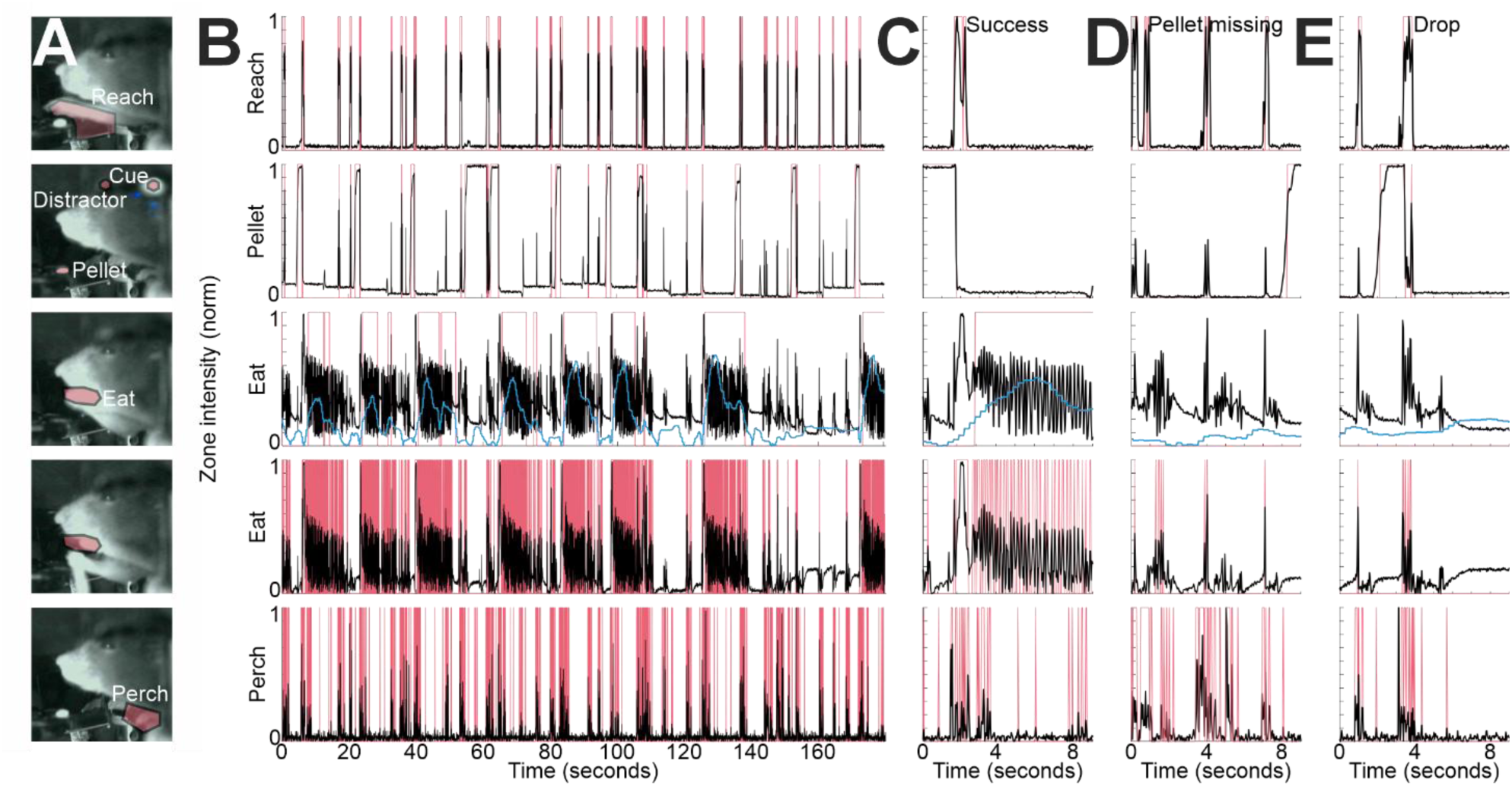
Regions of interest (“zones”) used for reach detection and outcome classification. **(A)** Positions of user-defined zones overlaid on frames from the video. **(B)** From a representative video segment during behavior, intensity signals within the zones normalized between 0 and 1 (black) and zone-specific event detection (red). Reach zone: Red when Z-scored signal exceeds a threshold (putative reach). Pellet zone: Red when signal exceeds a threshold defined by fitting a two-mode Gaussian mixture model to the signal (putative pellet available to mouse). Eat zone (upper row): Red when power of the signal between 5.5-7.7 Hz (light blue) exceeds a threshold (putative chewing). Eat zone (bottom row): Red when Z-scored signal exceeds an intensity threshold (putative entry of paw/pellet into region near mouth). Perch zone: Red when temporal derivative of Z-scored signal exceeds a threshold (putative movement including initiation of movement preceding reach). Close-up of signals across zones during an example successful reach **(C)**, reach when the pellet was missing **(D)**, and reach when mouse dropped the pellet **(E)**.

Specifically, we defined these ROIs as follows:

1. Reach Zone: The space between the initial resting position of the paw and the accessible pellet; the forelimb was required to enter this zone to retrieve the food.
2. Pellet Zone: The specific location of the accessible food pellet.
3. Eat Zone: The area surrounding the mouse’s jaw where the pellet arrived and where jaw oscillations occurred during chewing.
4. Perch/Paw Resting Zone: The resting position of the paws prior to reach initiation.
5. Lick Zone: The small space immediately in front of the mouth where the tongue protruded during licking behavior.

### Temporal Synchronization and IR Signaling

To synchronize Arduino-controlled behavioral events with the video frames, we positioned infrared (IR) LEDs directly against the camera face. Each LED was controlled by a dedicated output from the Arduino corresponding to specific events, such as the delivery of a cue. Consequently, whenever a cue was triggered, the camera recorded a set of illuminated pixels within a designated ROI. We utilized IR LEDs because mice cannot perceive IR light; furthermore, we shielded the rear of each LED to prevent any visible light leakage or reflections from reaching the animal. These LEDs functioned as additional behavior-relevant ROIs and were tracked by the analysis software to provide precise temporal alignment between hardware triggers and video data.

### Identification of Behavior-Relevant ROIs

To identify the behavioral ROIs, or zones (**Fig. 3A**), we used a custom MATLAB graphical user interface (GUI) that allowed a user to review video segments and manually define polygonal ROIs on a representative frame. While this manual approach was highly effective and provided precise spatial definitions, it required approximately 1-2 minutes of user intervention per video.

### Signal Extraction from ROIs

We extracted temporal signals from each ROI across every frame of a training session. These signals consisted of either the individual color channel intensities (red, green, and blue) or the overall mean pixel intensity derived from the video data.

### Heuristic Signal Processing and Event Classification

User-defined parameters for signal processing were managed via a configuration settings file. We implemented both manual and automated versions of the analysis; while the automated pipeline proved highly effective, we manually verified outputs and adjusted parameters whenever we altered the physical configuration of the behavior rig or to confirm data quality.

### Reach Detection

Reach events were identified by tracking pixel intensity within the reach ROI. Under infrared (IR) illumination, the forelimb appeared significantly lighter than the background, allowing forelimb entry into this zone to be detected as a signal increase (**Fig. 3B**, top row). We Z-scored the intensity data and identified reaches as discrete peaks exceeding a threshold of 2.5. To maintain physiological relevance, reaches were required to be separated by at least 166 ms (a maximum frequency of 6 Hz). Reach duration was defined by the start and end of the threshold crossing, with any reach exceeding five seconds classified as a “hold.” Occasionally, mice extended the forelimb and maintained contact with the surface of the pellet positioner disk for a prolonged period–a “hold.” We classified cases in which the forelimb entered the reach zone during chewing as “reaches,” but later discarded them if they (1) overlapped with an active chewing window (see below) or (2) occurred immediately after a successful reach, hence during chewing.

### Pellet Detection

Pellet availability was quantified using the intensity signal from the pellet ROI. High intensity signaled pellet presence, while low intensity indicated pellet absence. Users could choose either a manually user-defined threshold or an automated approach to determine the threshold for pellet presence. For the automated approach, we fit a two-mode Gaussian mixture model and used the mean of the higher mode as the threshold for “high” (**Fig. 3B**, second row).

### Chewing Detection

Because chewing-related jaw movements produced strong power in the 5.5-7.7 Hz frequency range in our videos, we detected chewing by measuring power in this band using the Chronux toolbox^35^. Behavior was classified as “chewing” when the Z-scored power exceeded 0.25 (**Fig. 3B**, third row).

### Paw Lifted to the Mouth Detection

We monitored the eat zone for a signal deflection (1.5 standard deviations above the Z-scored mean) within one second of a reach (**Fig. 3B**, fourth row). This deflection served as an indicator that the paw was successfully raised to the mouth. The absence of this signal caused any reach to be classified as unsuccessful.

### Fidgeting and Reach Initiation

To identify any paw movements prior to the forelimb extension, we monitored the perch zone. Fidgeting was defined as a frame-to-frame intensity change exceeding 2 standard deviations above the Z-scored mean (**Fig. 3B**, fifth row). This derivative-based approach allowed for clear separation between quiescent states and paw movement. Reaches were classified as one of two types: either beginning from the paw resting perch or not. Reaches were classified as beginning from the paw resting perch only if the animal remained at rest (zero fidgeting) for at least 0.5 seconds prior to the reach.

### Preliminary Success Classification

In the initial analysis pass, reaches were classified as successful (i.e., mouse consumed the pellet) or unsuccessful based on various temporal criteria (**Fig. 3C,D,E**): Pre-reach Chewing: Reaches were classified as unsuccessful if chewing activity was detected within the four-second window immediately preceding the reach. This was useful, because the experimental design used a minimum inter-pellet interval of nine seconds. Hence, any chewing from a prior successful trial was most likely complete before the next pellet became available. Delayed Initiation: Reaches were classified as unsuccessful if chewing failed to initiate within seven seconds post-reach. Insufficient Duration: Reaches were classified as unsuccessful if the total chewing time was less than six seconds within a 20-second post-reach window.

### Random Forest Classifier to Discriminate Successes and Drops

To further improve the discrimination of successful and unsuccessful reaches when the mouse dropped the pellet, we trained a Random Forest Classifier to separate these reach outcomes. First, we applied Otsu’s method^36^ to the chewing power signal to differentiate between chewing and non-chewing states. For each reach attempt, we analyzed a 1.667-second window (50 frames) starting from reach onset and extracted three primary features: average chewing power (5.5-7.7 Hz), chewing duration, and peak pixel intensity within the eat zone. These features were paired with manually annotated ground-truth labels to assemble a dataset of 19140 reach events across 345 sessions. We used a stratified 80/20 split by reach event, yielding 15312 training events and 3828 testing events.

### Checking the Automated Analysis

We validated the automated output by cross-referencing classified events with raw video frames. Using a custom function (checkBehaviorOutput.m), we printed the movie frame indices associated with various event types, then manually verified classifications by viewing the raw video in Fiji^37^. We performed these audits on a representative data subset and whenever we modified the behavioral rig or behavior paradigm.

### Hardware Synchronization and Data Export

To reconcile the independent reference frames of the Arduino and the video, we used a “random” IR LED signal for temporal alignment. This allowed us to synchronize events recorded by the Arduino (such as internal pellet loading) with the video data. Cue onsets were detected directly from the video via a dedicated cue ROI tied to the Arduino-controlled IR LED. Final data were exported both as continuous temporal vectors for each event type and as trial-aligned matrices centered on cue onset.

### Testing DeepEthogram

To compare our heuristic pipeline with a published deep-learning method called DeepEthogram^38^, we applied DeepEthogram to 1126 segments from the behavioral videos. The data were partitioned into an 80% training set and a 20% validation set. We trained DeepEthogram to identify five discrete classes: no reaching, successful retrieval, pellet drop, failure to dislodge the pellet (called a miss), and reach when the pellet was absent. We measured the accuracy using the validation set.

### Measuring Reach Cancellation

Reach progression was defined by six stereotypical, sequential phases: initial reach, full extension, pellet grasp, retrieval, paw-to-mouth, and return-to-perch. A reach was classified as “cancelled” if the mouse deviated from this standardized sequence or abandoned the attempt prematurely. Temporal onset was marked at the moment of deviation. Notably, a reach was only deemed cancelled if the behavioral sequence was disrupted; failure to retrieve or hold the pellet did not constitute cancellation if the motor program reached completion. Cancellation was identified by phase-specific signatures, such as the cessation of forward movement, loss of pellet contact, or trajectory deviations. These were further corroborated by subtle behavioral indicators, including atypical facial movements, aberrant limb positioning, or postural shifts, which often served as early predictors of intent to abandon the sequence.

### Measuring the Cancellation Time: Drops

We automatically detected drops using the analysis pipeline. Then we manually determined the cancellation time by watching video segments centered on drops.

## Results

### Pellet Loading and Positioning

We evaluated the mechanical performance of the behavior system by measuring its pellet presentation spatial accuracy. The system delivered pellets with diameter 3.175 mm to a consistent location ±3 mm >90% of the time (**Fig. 4**). This ensured that the reach target remained in a predictable location for the mouse during most reaching attempts.

**Figure 4.**
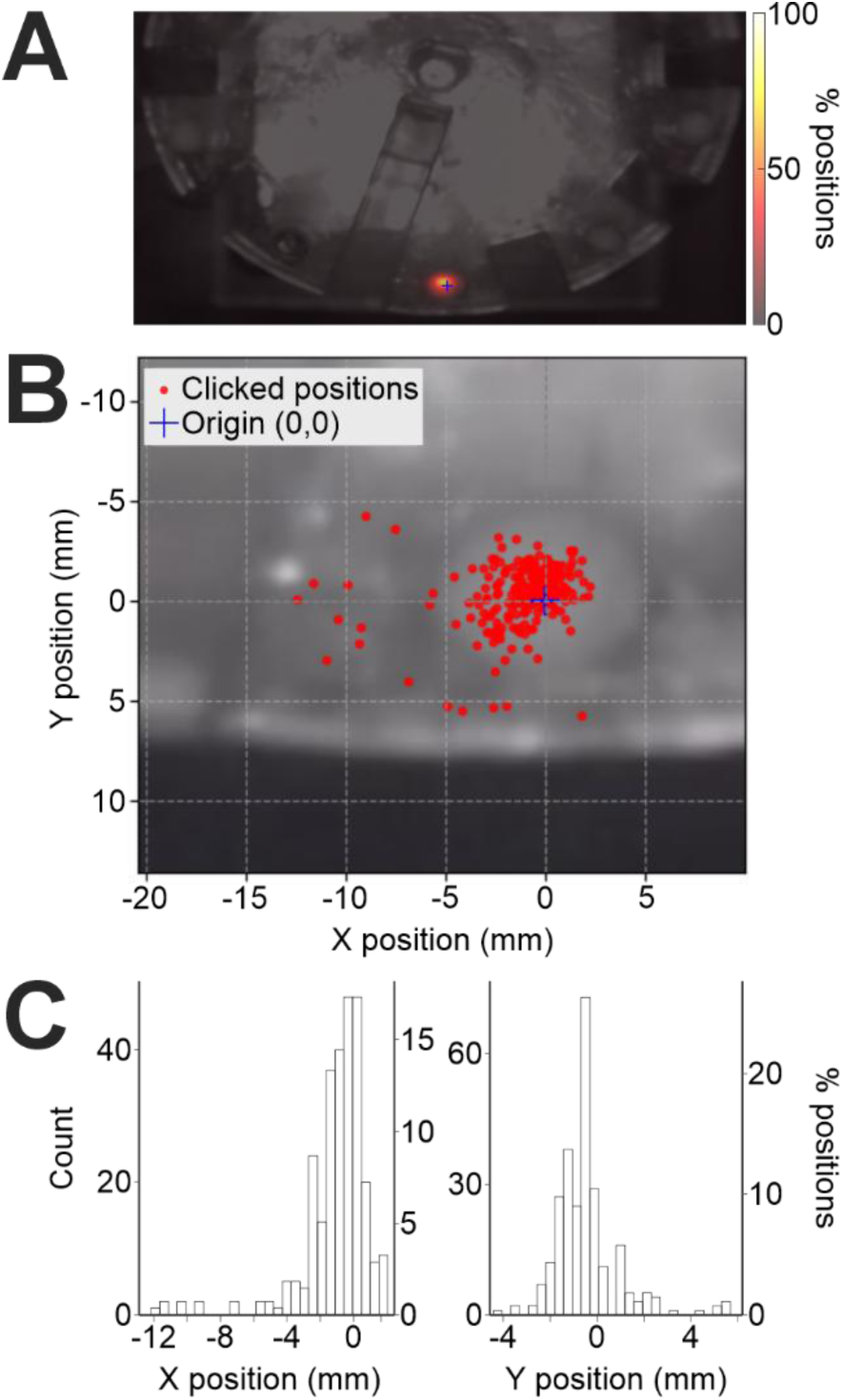
Consistency of pellet positioning by automated rig. **(A)** Gaussian-smoothed heat map of pellet positions overlaid on image of pellet positioner disk (276 loading attempts). **(B)** Example image showing pellet positions logged in X and Y. **(C)** Histograms showing pellet positioning accuracy in X and Y.

### Behavioral Training Success Rate

We assessed the efficiency of our mouse training protocol by calculating the fraction of mice that successfully mastered the reaching task. Out of 111 mice trained over a 15-day period, 97 (87.4%) met the designated success criterion, as previously reported in Reinhold et al.^39^. This criterion was the ability to retrieve and consume 20 or more pellets within a single one-hour session. In contrast, less than 10% of head-fixed mice produced reaches spontaneously (without experimenter-led behavioral shaping) within the first 3 days on the rig.

### Classification Accuracy and Performance Comparison

We quantified the accuracy of the automated pipeline by comparing its outputs to reach classifications by a human (**Fig. 5A**). The no reach detection accuracy (i.e., binary classification as not reaching or reaching) was 92%. We then considered whether the automated classification code could determine the outcome of a reach. The code correctly identified 96% of successful reaches, 91% of dropped pellets, 98% of attempts where the pellet was not dislodged from the loading hole, called misses, and 85% of reaches when no pellet was available.

**Figure 5.**
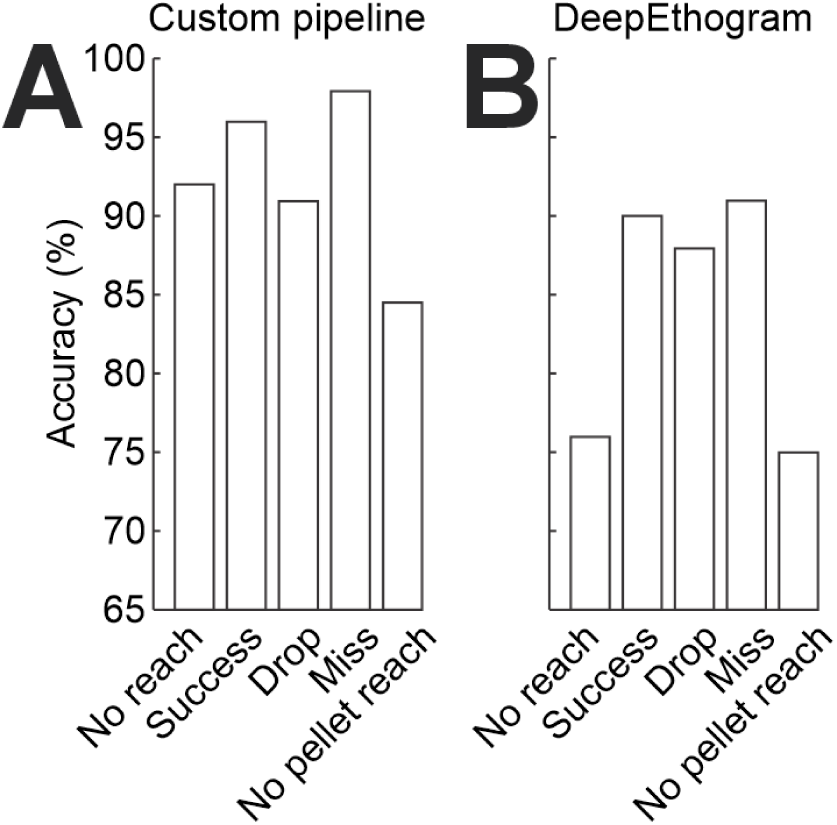
Accuracy of reach detection and outcome classification pipeline. **(A)** Accuracy of custom detection pipeline compared to manual annotation. Of 75 pipeline-detected reach events, 69 were identified by a human as forelimb extensions (92%), and 6 were oral-motor movements (chewing). Classification accuracy of reach outcomes: successful (103/107, 96%), dropped pellets (86/94, 91%), and missed attempts (89/91, 98%). The accuracy for classifying reaches when no pellet was available was 85% (87/103). A subset of the data in this panel was previously reported^39^ and is replotted here to illustrate performance. **(B)** Performance of DeepEthogram classifier compared to manual annotation across 225 video segments. 171 out of 225 (76%) were real forelimb extensions toward the pellet. Classification accuracy of reach outcomes: successful (202/225, 90%), dropped pellets (198/225, 88%), and missed attempts (205/225, 91%). The accuracy for classifying reaches when no pellet was available was 75% (169/225).

In comparison, analysis using DeepEthogram (**Fig. 5B**) yielded accuracies of 90% for successful reaches, 88% for dropped pellets, 91% for misses, and 75% accuracy for reaches when no pellet was available. Hence, overall DeepEthogram was successful at classifying the outcome of the reach. The no reach detection accuracy (called “Background” in DeepEthogram) was 76%, although further training might have improved this number. Hence, our custom pipeline showed slightly higher accuracy rates, although DeepEthogram may be a more plug-and-play approach requiring less manual tuning of parameters.

### Reach Phases

We quantified the duration of each phase of the reach (**Fig. 6**). Across mice, the initial ballistic extension of the forelimb (∼150 ms on average) was faster than the later movement lifting the pellet toward the mouth (∼280 ms on average).

**Figure 6.**
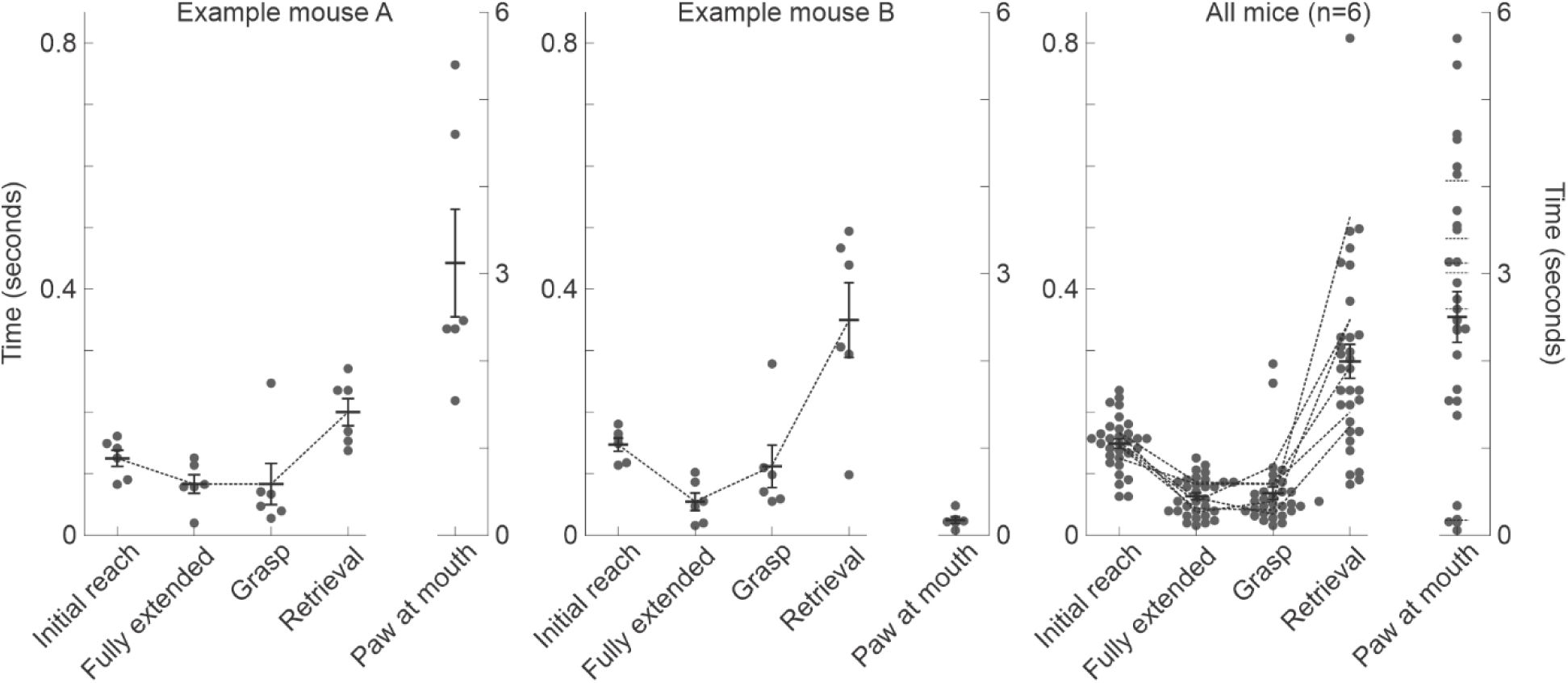
Duration of reach phases. Reach phase durations based on high-speed video annotation. Reach phases defined as follows. Initial reach: from paw first lifted to fully extended forelimb. Fully extended: time spent with forelimb fully extended as paw hovers over pellet. Grasp: from paw touching the pellet to first lifting the pellet. Retrieval: from the paw first lifting up the pellet to the pellet first touching the tongue. Paw at mouth: from initial tongue contact to initiation of paw withdrawal. Each point is a reach. Mean and s.e.m. are the summary statistics. (Left two columns) Representative examples from two mice illustrating inter-subject consistency across the first four reach phases, contrasted with variability in the final phase. The terminal phase (paw at mouth) during chewing revealed individual-specific movement signatures: Mouse A typically maintains the paw near the mouth during chewing, whereas Mouse B withdraws the paw to the starting perch. (Right column) Summary across six mice. Each dotted line is one mouse.

### Demonstration of Scientific Utility: Measuring Reach Cancellation Time

To demonstrate the sensitivity of the behavioral pipeline, we examined interruption of an ongoing reach. We measured the minimum time required for the mouse to initiate canceling a reach after dropping the pellet. This was not instantaneous. We focused on the minimum cancellation time rather than the mean time to place a lower bound on the speed of initiating cancellation. The physical slippage of the pellet out of the paw served as the cue to cancel the reach. We measured the delay between slippage and the first clear sign of reach cancellation. We measured this cancellation time, which included the time to detect pellet slippage, as a function of three reach phases: (1) fully extended/initial grasp, (2) retrieval, or (3) pellet entry into the mouth. The minimum cancellation time (MCT), defined as the 10th percentile of the distribution of cancellation times, varied across reach phases, increasing from 120 ms during the initial grasp to 180 ms at pellet entry into the mouth (**Fig. 7**, permutation test p-value comparing these MCTs: p=0.03). Thus, the time required to initiate cancellation was longer than the physical duration of individual reach components, such as arm extension and grasping (**Fig. 6**). Furthermore, this cancellation time increased as the reach neared completion and the pellet approached the mouth.

**Figure 7.**
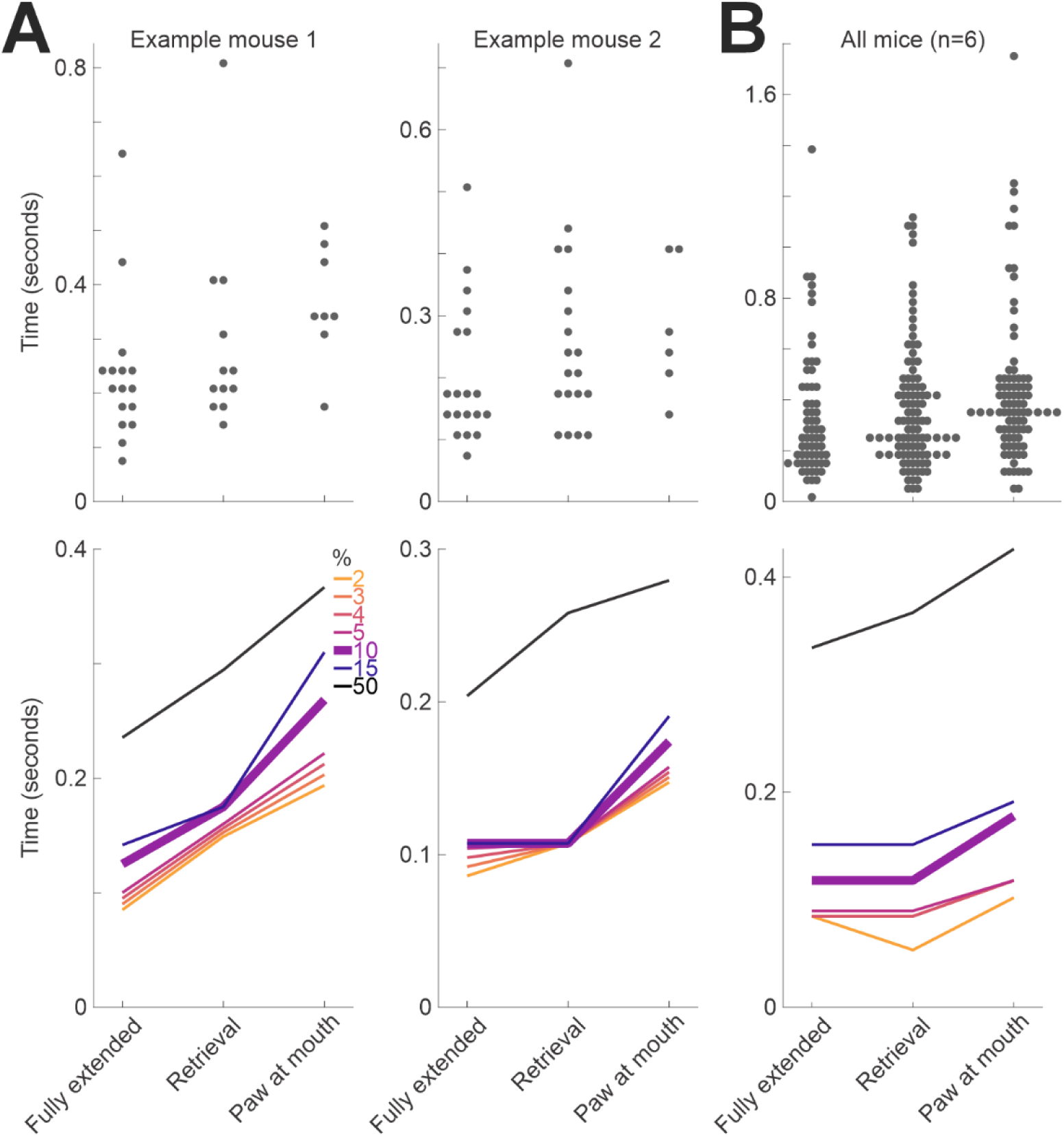
Minimum cancellation time as a function of reach phase. **(A)** Minimum time required for the mouse to initiate cancellation of the reach after the pellet was dropped across two representative example mice. Top row: all analyzed drops. Bottom row: summary percentiles including the minimum cancellation time (MCT), defined as the 10th percentile of the distribution, in purple. **(B)** Across 6 mice. Layout as in A. One-sided permutation tests assessed whether the MCT increased between fully extended and paw at mouth, or showed an increasing linear trend across reach phases. Null distributions were generated by shuffling observations across groups while preserving group sizes. Fully extended vs. paw at mouth: p = 0.0300; slope across phases: p = 0.0599.

## Discussion

Head-fixed reaching in mice is increasingly being used in neuroscience to investigate skilled motor control. We have presented a comprehensive protocol for this paradigm with the aim of assisting the community in implementation. Specifically, we have provided: (1) a do-it-yourself (DIY) behavior rig design that is simple and reproducible; (2) an effective behavioral shaping method for training mice to perform skilled reaches; (3) automated analysis code for video-based classification; and (4) a demonstration of the scientific utility of this approach.

### Behavior Rig

The DIY rig, unlike many other currently available open-source systems, was designed to precisely position the food pellet in front of the mouse. The rig was open-loop but accurate at pellet positioning (90% spatial accuracy to within 3 mm), though pellet orientation was not controlled. Although <100%, we found that this loading accuracy was sufficient for our experimental needs, because the analysis code used the video to explicitly verify successful pellet delivery in every case.

### Training

Our shaping protocol for reach training resulted in a high learning rate (87.4%). Without this shaping protocol, only a small fraction of mice were able to generate reaches spontaneously in the head-fixed position. We observed that certain phases of the reach were easier to acquire than others. The initial ballistic extension of the forelimb required minimal training. In contrast, the retrieval phases of grasping, supination, and raising the pellet to the mouth required more practice. These retrieval phases were also slower, perhaps suggesting that the later phases of the head-fixed reach might be particularly dependent on motor learning.

### Advantages and Disadvantages of the Behavioral Paradigm

This behavioral paradigm has several advantages. The task enables the study of motor control, and complete-trajectory paw tracking is simpler than complete-trajectory tongue tracking, as the tongue is still inside the mouth at lick initiation. Reaching toward a distant pellet is relatively slower than licking, and hence the task also introduces a long inherent delay (∼150 ms) between cue and reward. In addition, failures (drops) persist even in expert mice, providing a useful window into unsuccessful trials. However, the paradigm also has limitations. It requires a manual training stage, depends on the animal’s fine motor control, and involves the unnatural constraint of head fixation.

### Automated Analysis Code

Our analysis pipeline was designed as an “expert system” tailored to our specific task, achieving high classification accuracy at detecting reaches and discriminating reach outcomes (between 85% and 98% depending on reach type). Compared to DeepEthogram, our approach was computationally less intensive, but it required user input (drawing ROIs). Presently this user input (∼2 minutes per video) is the most manual component of the analysis and an area for future improvement. In practice, however, these ROIs were often stable across days, reducing the frequency with which this step must be repeated. Our expert system works, because fundamentally the signals indicating different outcomes were clear and highly interpretable.

However, DeepEthogram may offer a more plug-and-play solution for some users. Perhaps, a future combination of simple ROI-based reach detection and deep learning-based outcome classifications would be optimal.

### Scientific Demonstration

We also used this paradigm to examine interruption of an ongoing reach as a proof of principle for the temporal sensitivity of the behavioral pipeline. We made two observations. First, the minimum cancellation time varied across phases of the reach. Second, the minimum cancellation time increased later in the movement. In the later phase of the movement, the cancellation time was often longer than 180 ms. This long delay hints at a cognitive decision process beyond a pure motor constraint. However, multiple factors could contribute to this phase dependence, including differences in movement kinematics, motor complexity, detection time differences, or sequence structure.

In conclusion, this head-fixed reaching paradigm represents a rich, well-controlled behavior for studying the mechanisms of learned motor sequencing and cancellation.

## Author Contributions and Acknowledgments

A.C. and K.R. designed the behavioral rig. K.R., A.M., and J.W. wrote the analysis code. K.R. developed the training protocol. J.B.W. performed testing of the DeepEthogram tool. K.R. wrote the manuscript. We thank Bernardo Sabatini for his support and guidance.

## Competing Interests

The authors declare no competing interests.

## Funding

Preparation of this manuscript was supported in part by the National Institutes of Health under Award Number R00MH127471. Development of the pipeline was supported by institutional and laboratory funds during K.R.’s postdoctoral training.

## Data Availability

Data will be made available from the corresponding author upon reasonable request.

## Code and Materials Availability

The Reinhold Lab website will serve as the primary distribution page for updated designs, analysis code, Arduino control code, CAD files, Illustrator files, assembly materials, and related resources: https://www.reinholdlab.org/resources. Current code versions are available at: https://github.com/kimerein/reach-behavior-analysis,

https://github.com/kimerein/behaviorRig, https://github.com/kimerein/reaching-behavior-rig. Designs and materials are also available from the corresponding author upon reasonable request.

## Notes

https://www.reinholdlab.org/resources

### Competing Interest Statement

The authors have declared no competing interest.

